# Oxidative stress is a potential cost of synchronous nesting in olive ridley sea turtles

**DOI:** 10.1101/2021.06.21.449061

**Authors:** B. Gabriela Arango, Martha Harfush-Meléndez, David C. Ensminger, Elpidio Marcelino López-Reyes, José Alejandro Marmolejo-Valencia, Horacio Merchant-Larios, Daniel E. Crocker, José Pablo Vázquez-Medina

## Abstract

Olive ridley sea turtles, *Lepidochelys olivacea*, exhibit a polymorphic reproductive behaviour nesting in solitary or in mass aggregations termed “arribadas”, where thousands of turtles nest at once. Arribadas may provide fitness benefits including mate finding during nearshore aggregations and predator satiation at the time of hatching, but the potential costs of arribada nesting remain understudied. To explore the potential trade-offs of the fitness benefits associated with arribada nesting, we collected blood from olive ridley turtles nesting in arribada and solitary. We measured reproductive and metabolic hormones (progesterone, estradiol, testosterone, thyroxine, and triiodothyronine), triglycerides (TG), non-esterified fatty acids (NEFA) and markers of oxidative damage (4-hydroxynonenal, malondialdehyde, protein carbonyls, and nitrotyrosine). Arribada nesters were bigger and had higher levels of progesterone than solitary nesters. Similarly, thyroid hormones were higher in individuals nesting in arribada than in solitary nesters, while TG and NEFA were positively correlated in arribada but not in solitary nesters. Nesting in arribada was associated with increased lipid peroxidation and protein carbonyls compared to solitary nesting. These results suggest that nesting in arribada is potentially more energetically expensive than nesting solitarily, and that oxidative stress may be a trade-off of the fitness benefits associated with arribada nesting.

## Introduction

Sea turtles of the genus *Lepidochelys*, which includes Kemp’s (*Lepidochelys kempii*) and olive ridley turtles (*Lepidochelys olivacea*), exhibit an unusual reproductive polymorphic behaviour, nesting both in solitary and in massive aggregations termed “arribadas” [1]. During arribada nesting, thousands of turtles nest synchronously over a 2-7 day interval [1–3]. Notably, the same individual can nest interchangeably in arribadas or solitary, but the factors that determine whether an individual joins the arribada or nests in solitary remain unknown [2,4].

Olive ridley turtles nesting in arribadas can extend their inter-nesting intervals up to 63 days by inducing hypoxia-driven pre-ovipositional embryonic arrest so they can synchronize to nest [5,6]. Environmental cues that might trigger arribadas include the moon phase, weather patterns, and rainfall [2,7]. Although the exact benefits of arribada nesting are unknown, this behaviour provides a social context that promotes multiple mating and paternity, increasing genetic exchange by three-fold compared to solitary nesting [2,8–10]. Furthermore, nesting in arribada increases early hatchling dispersal by overwhelming or satiating predators [11]; however, arribada nesting does not necessarily promote hatchling success as increased organic matter from egg saturation and high nesting density in arribada beaches result in lower hatchling success compared to solitary nesting [3,12,13]. In contrast, solitary nesters produce bigger hatchlings, which have higher male-to-female ratio than hatchlings produced by arribada nesters [1,6]. Thus, while both nesting modes are likely important for the population, it is unknown if the potential fitness benefits associated with arribada nesting come with a physiological cost.

A central principle of life history theory is that reproductive effort negatively affects survival; however, there is conflicting evidence about the proximal costs of reproduction [14]. One of the leading hypotheses suggests that oxidative stress is a potential trade-off for reproductive investment [15–17], but other studies do not support this idea [18,19]. Work in wild vertebrates suggests that diversification of reproductive strategies is associated with sexual differences in oxidative stress [20–22]. In male and female free-ranging macaques [23], male northern elephant seals [24] and female North American red squirrels [25], reproduction causes oxidative stress. Similarly, nesting decreases the activity of the antioxidant superoxide dismutase in female loggerhead sea turtles [26].

To the best of our knowledge, the potential trade-offs of arribada nesting have not been studied. Therefore, we compared the circulating concentrations of reproductive and metabolic hormones, and markers of oxidative stress in olive ridley turtles nesting solitary and in arribada. We found that arribada nesters are heavier and have higher metabolic activity and oxidative damage than solitary nesters. Thus, our results suggest that oxidative stress is a potential cost of arribada nesting in olive ridley sea turtles.

## Methods

### Animals and sample collection

Animal handling protocols were approved by Sonoma State University’s IACUC. Samples were collected under permit SGPA/DGVS/12915/16 and imported to the US under permits CITES MX88143 and CITES 19US85728C/9. Nesting olive ridley turtles were sampled at the marine protected area of *La Escobilla*, Oaxaca, Mexico (15° 47’ N; 96° 44’ W) during arribada (n = 13). Solitary nesting turtles were sampled at *Campamento Tortuguero Palmarito*, Puerto Escondido, Oaxaca, México (15° 53’ 26.3” N; 97° 07’ 52.2” W), and *La Escobilla* during solitary nesting (n = 10). Samples were collected after the animals had dug their nest, during the ‘trance-nesting period’ [27]. None of the animals were disturbed from nesting, nor returned to the ocean without laying their eggs. Mass was estimated using a regression from published olive ridley morphometric data (n= 59, mass = −47.44 + 1.13 x straight carapace length (SCL), r^2^ = 0.70, p < 0.001; [28]). Animals sampled during solitary nesting were weighed using a hand-held scale (± 0.1 kg) to assess the validity of the mass-estimation method. The equation used to estimate mass predicted the weight of solitary nesters with a mean error of 4%. Blood was collected from the cervical vein into chilled heparin Vacutainer tubes [27,29]. Plasma was prepared by centrifugation onsite, frozen, and subsequently stored at −20°C until laboratory analysis.

### Biochemical Assays

#### Hormones

Progesterone (P4), estradiol (E2), thyroxine (T4), triiodothyronine (T3) and testosterone were measured in sea turtle serum using ELISA or RIA kits (progesterone: MP Biomedical ELISA catalog number 07BC1113, estradiol: MP Biomedical ELISA catalog number 07BC1111, T3: MP Biomedical RIA catalog number 06B-254215, T4: MP Biomedical RIA catalog number 06B-254011, testosterone: Enzo ELISA catalog number ADI-900-065). Assay platforms were validated for use in olive ridley turtles. Serially diluted pooled samples (1:2 to 1:16) exhibited parallelism to the standard curve after log-logit transformation. Mean recovery of hormone added to serum pools was 101.5 ± 3.5%, 98.3 ± 6.5%, 99.4 ± 4.7%, and for 101.3 ± 5.1% (r^2^ > 0.98) for P4, E2, T3 and T4, respectively. The testosterone kit was previously validated for sea turtle blood [30] and was further validated for our samples using parallelism. Assays were conducted following manufacturer’s instructions.

#### Oxidative damage

Two markers of lipid peroxidation (4-hydroxynonenal: 4-HNE, malondialdehyde: MDA), a marker of protein oxidation (protein carbonyls), and a marker of protein nitration (3-nitrotyrosine) were measured in sea turtle plasma using ELISA kits (Cell Biolabs catalog numbers: STA-838, STA-832, STA-310, STA-305). Assays were conducted following the manufacturer’s instructions with minor modifications as previously described [31].

#### Plasma lipids and total protein

Plasma triglycerides (TG) and non-esterified fatty acids (NEFA) were measured using colorimetric kits (TG: Cayman Chemical catalog number 10010303, NEFA: Wako Chemicals, HR Series NEFA-HR (2) catalog number: 999-34691). Total protein content was measured using a bicinchoninic acid (BCA) protein assay (Thermo Fisher Scientific catalog number: A53227). Oxidative damage values were normalized to total plasma protein levels.

All samples were analyzed in duplicate in a single assay. The average intra-assay coefficient of variation was <6%.

### Statistical analyses

Statistical analyses were conducted using JMP Pro 15 (SAS Institute, USA). Equality of variances was assessed using Levene’s test. Progesterone, TG and NEFA data were log10 transformed to meet model assumptions. Arribada and solitary nesting groups were compared using two-sample t-tests. A nominal logistic regression was used to analyze 4-HNE data since the response variable was binary. Correlation analyses between hormones, mass, and metabolites including those measured in samples from the same animals in a previous study [32], were conducted by calculating Pearson or Spearman coefficients (for corticosterone non-parametric data). Data are expressed as mean ± SD unless otherwise indicated. Statistical significance was considered at p≤0.05. Figures were generated using R packages ggplot2, plotly, scales and ggthemes.

## Results

### Arribada nesters have higher progesterone levels than solitary nesters

We measured circulating progesterone, estradiol, and testosterone to assess reproductive hormone profiles in solitary and arribada nesters. Progesterone was higher in arribada than in solitary nesters (3.58 ± 1.71 vs 2.10 ± 1.56 ng/mL, t = 2.612, p = 0.02; **Figure 1A**). Estradiol did not differ between nesting conditions (solitary 4.47 ± 2.79 vs arribada 5.7 ± 2.24 pg/mL, t=-1.17, p=0.25; **Figure 1B**). Similarly, testosterone was not different between solitary and arribada nesters (70.43 ± 42.88 vs 75.08 ± 35.99 pg/mL, t=-0.29, p=0.78; **Figure 1C**). These results show that females nesting in arribadas have higher progesterone but not estradiol or testosterone levels than solitary nesters.

**Figure 1.**
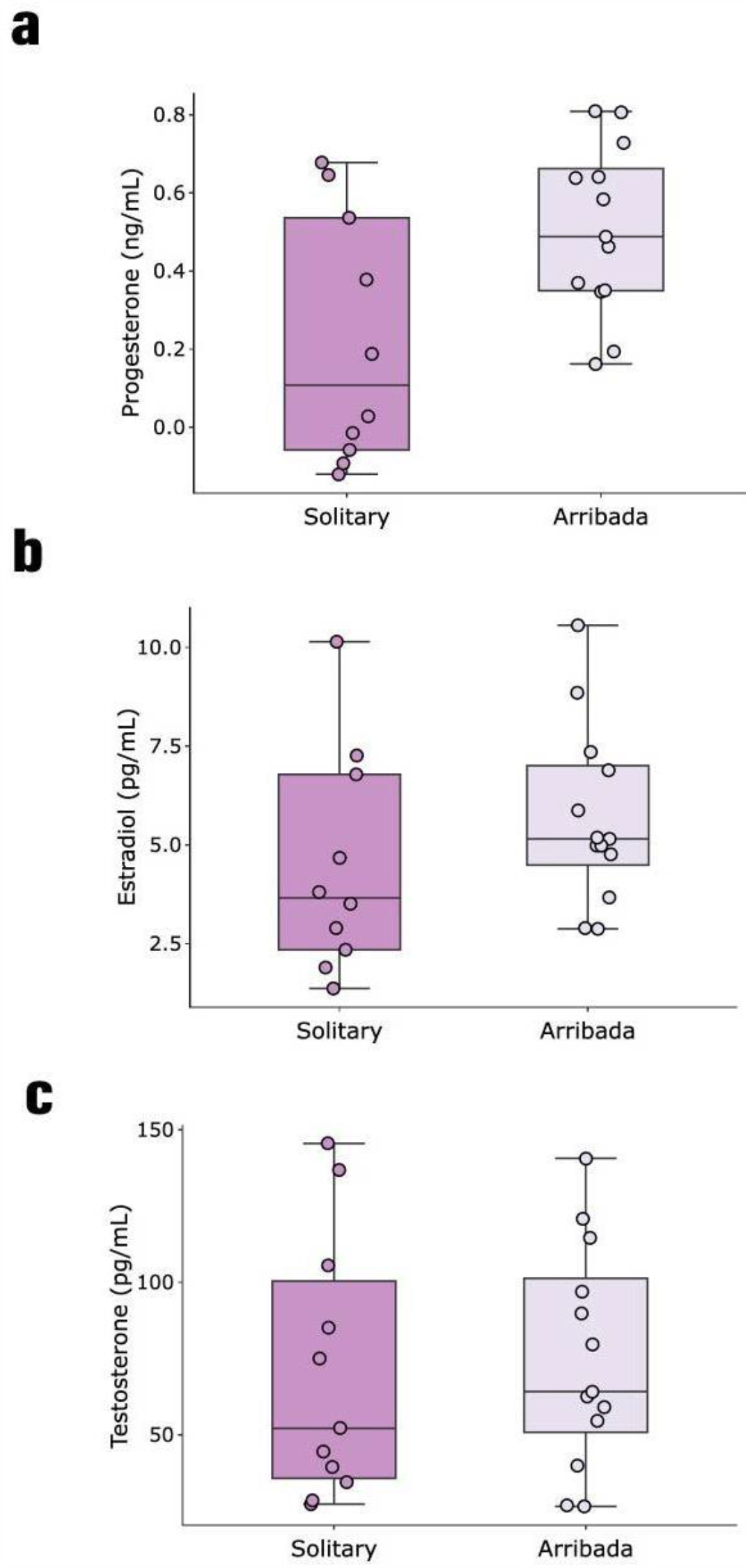
Progesterone levels are higher in arribada than in solitary nesters. Reproductive hormone levels in solitary and arribada nesters. **A)** Progesterone (p = 0.02), **B)** estradiol (p = 0.25), and **C)** testosterone concentrations (p = 0.78).

### Metabolic activity is higher in arribada than in solitary nesters

We quantified plasma T4 and T3 to evaluate whether arribada nesting induces a potential change in metabolic activity compared to solitary nesting. Both T3 and T4 were higher in individuals nesting in arribada than in solitary nesters (T3: 48 ± 19.54 vs 29.3 ± 11.86 ng/dL, t = 2.66, p = 0.015; T4: 1.45 ± 0.27 vs 1.12 ± 0.13 ug/dL, t = 3.62, p = 0.002; **Figure 2A, 2B**), suggesting increased thyroid function in arribada nesters. While neither TG nor NEFA levels were different between nesting modes (TG: solitary 14.46 ± 9.52 vs arribada 20.47 ± 10.03 mg/mL, t=1.94, p=0.07; NEFA: solitary 0.93 ± 0.22 vs arribada 1.08 ± 0.55 mM, t=0.445, p=0.66), arribada nesters were heavier than solitary nesters (29.20 ± 5.20 vs 24.76 ± 4.37 kg, t=2.17, p=0.042). We then conducted correlation analyses between hormones and metabolites including those (corticosterone, glucose and lactate) measured in samples from the same individuals in a previous study [32]. We found a positive correlation between corticosterone and glucose (ρ=0.87, p<0.0001; **Figure 3A**), corticosterone and lactate (ρ=0.73, p=0.005; **Figure 3B**), glucose and lactate (r=0.85, p=0.0003; **Figure 3C**), and TG and NEFA (r=0.82, p=0.0005; **Figure 3D**) in arribada but not in solitary nesters. These results suggest that arribada nesters have higher metabolic activity than solitary nesters.

**Figure 2.**
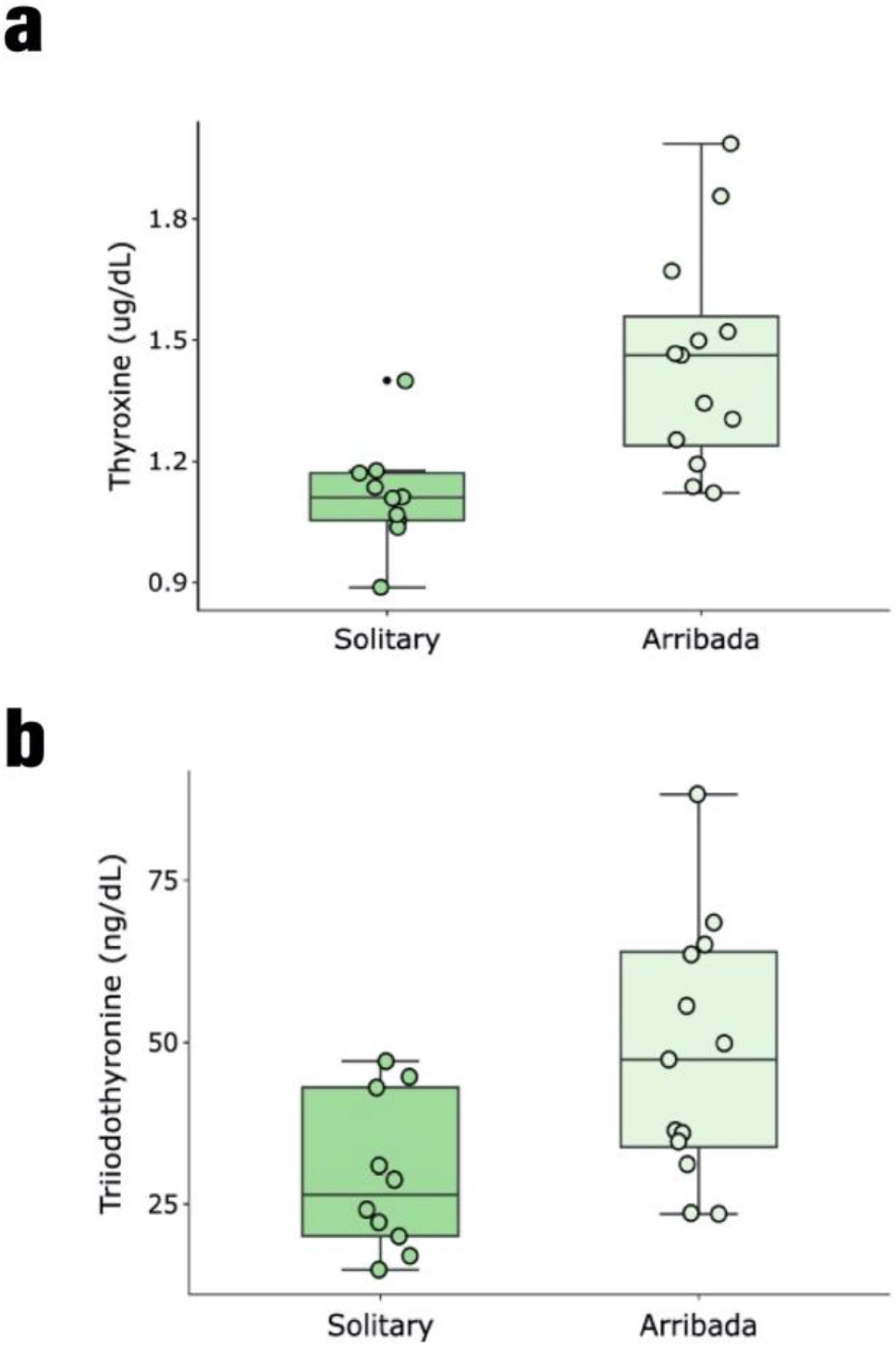
Thyroid hormones are higher in arribada than in solitary nesters. **A)** Thyroxine (p = 0.002) and **B)** triiodothyronine (p = 0.015) levels in solitary and arribada nesters.

**Figure 3.**
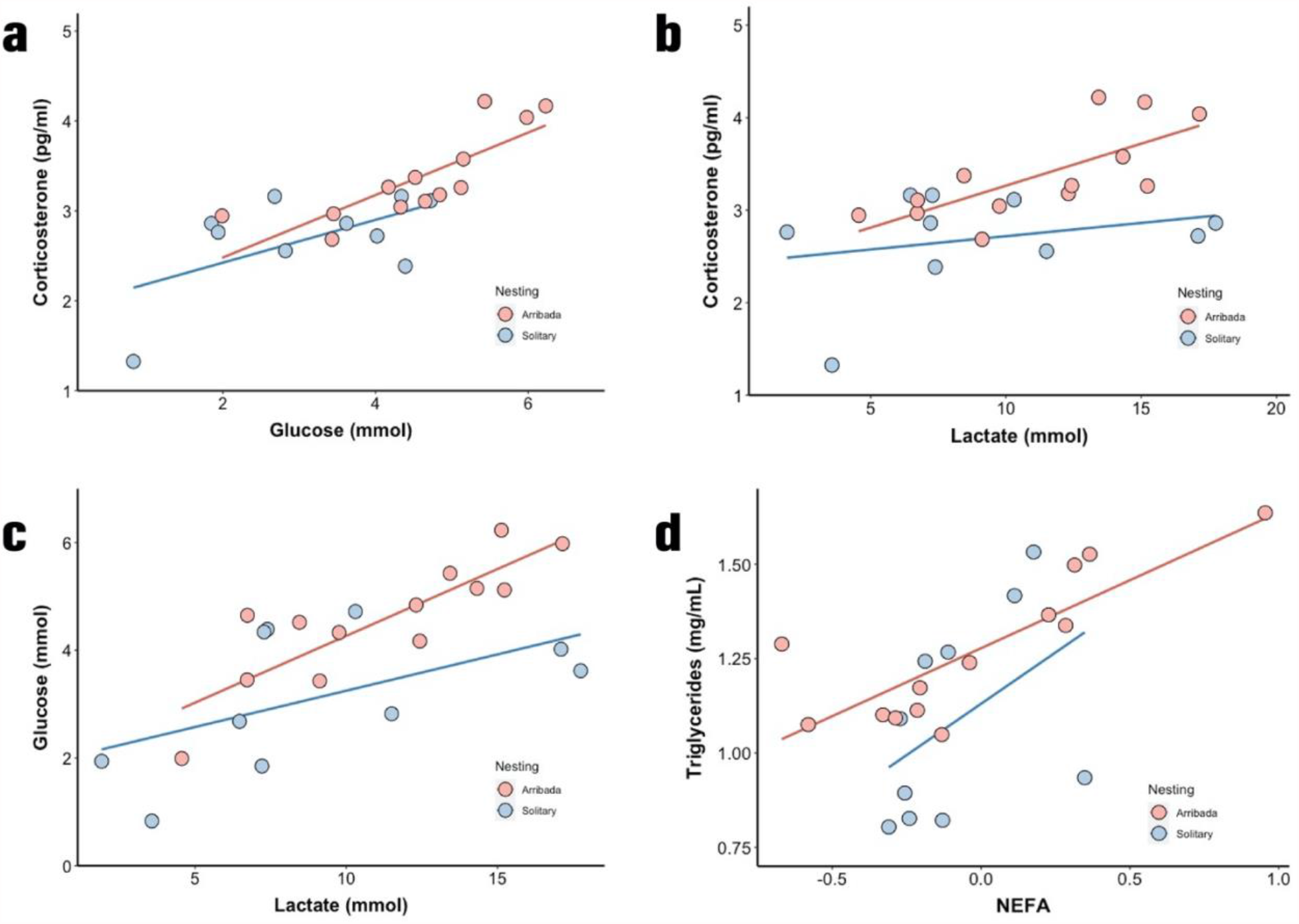
Correlations between hormones and metabolites in arribada and solitary nesters. **A)** corticosterone and glucose (arribada: ρ=0.87, p<0.0001), **B)** corticosterone and lactate (arribada: ρ=0.73, p=0.005), **C)** glucose and lactate (arribada: r=0.85, p=0.0003), **D)** TG and NEFA (arribada: r=0.61, p=0.0021).

### Oxidative damage is a potential cost of arribada nesting

We then evaluated whether oxidative damage represents a proximate cost for nesting in arribada by measuring four circulating markers of oxidative damage. Plasma levels of the lipid peroxidation products 4-HNE (arribada 100% positive vs solitary 25% positive for 4-HNE, **χ**^2^(2)=10.01, p=0.0016), and MDA (0.0025 ± 0.0013 vs 0.0011 ± 0.00034 pmol/***μ***g of protein, t=3.77, p=0.0019), along with protein carbonyls (0.00246 ± 0.0013 vs 0.00103 ± 0.00032 nmol/***μ***g of protein, t=3.64, p=0.0027) were higher in arribada than in solitary nesters (**Figures 4A, 4B, 4C**). In contrast, protein nitration (3-Nitrotyrosine) did not vary between nesting modes (solitary 0.039 ± 0.026 vs arribada 0.058 ± 0.031 nM/mg, t=-1.29, p=0.22; **Figure 4D**). These results suggest that oxidative damage is a potential cost of synchronous nesting in olive ridley sea turtles.

**Figure 4.**
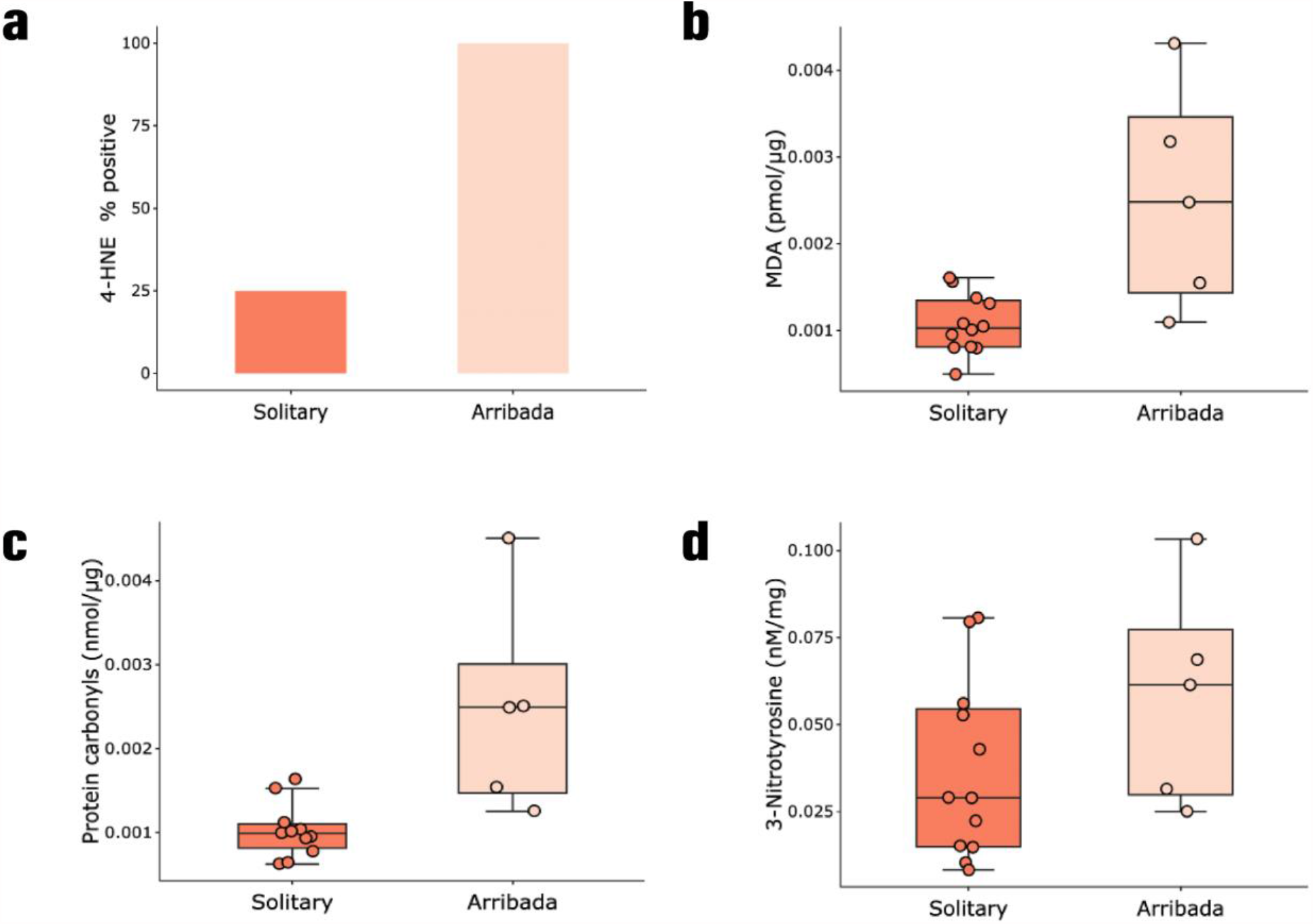
Oxidative damage is higher in arribada than in solitary nesters. Lipid peroxidation measured as **A)** 4-HNE-protein adducts (binary test, percent positive arribada, 100% vs solitary, 25%, p = 0.0009, **χ**2(2)=10.01, p=0.0016), and **B)** MDA-protein adducts (p = 0.0019); **C)** protein carbonyls (p = 0.0027) and **D)** 3-Nitrotyrosine (p = 0.22).

## Discussion

Every year thousands of olive ridley sea turtles synchronize in “arribadas” to nest over a 2-7 day period. The fitness benefits associated with arribada nesting include mate finding during nearshore aggregations [2], predator satiation at the time of hatchling [11], and multiple paternity and genetic exchange [8]. Whether arribada nesting carries a physiological cost remains unknown. Here, we found that arribada nesters have higher progesterone levels, metabolic activity and oxidative stress than solitary nesters. Therefore, our results suggest that nesting in arribada may be energetically more expensive than nesting in solitary and that oxidative damage is a trade-off for the potential fitness benefits associated with arribada nesting in olive ridley sea turtles.

### Potential endocrine drivers of arribada nesting

Environmental cues that trigger arribada events include the moon phase, temperature, humidity and salinity [2,7], but the exact combination of factors remains unknown and it is likely population-specific [7,33]. Similarly, endogenous adjustments that allow individuals to synchronize to join an arribada are poorly understood. Long-term egg retention is uniquely observed in the genus *Lepidochelys* spp. [5], where average inter-nesting intervals last about 28 days while inter-nesting in other species lasts 9-15 days [5]. Moreover, eggs from olive ridley turtles nesting in arribada can tolerate hypoxia-driven pre-ovipositional embryonic arrest for longer periods compared to eggs from solitary nesters (15 vs 4 days) [6]. Increased concentrations of gonadosteroids might also influence whether an individual joins the arribada. We found higher progesterone but not estradiol or testosterone levels in arribada than in solitary nesters. Previous work, however, shows that neither progesterone nor estradiol influence the holding of arribada eggs inside the reproductive tract [34], and that inter-nesting differences among *Lepidochelys* spp. and other sea turtles are not related to differences in ovulatory cycles, but rather to egg retention capacity [5,35]. A progesterone surge during ovulation induces a rapid albumen release into the oviduct, which activates previously stored sperm [36]. Therefore, rather than exerting endocrine control over synchronicity, it is possible that high progesterone levels are related to increased mating opportunities in arribada compared to solitary nesters. Of note, progesterone levels do not differ with nesting mode in Kemp’s ridley turtles [37]. Thus, there is no conclusive evidence about potential endocrine drivers of arribada nesting in olive ridley sea turtles.

### Is arribada nesting more energetically expensive than solitary nesting?

In a previous study conducted in the same individuals we showed that arribada nesters have higher circulating corticosterone and glucose levels than solitary nesters [32]. Here, we found that arribada nesters are bigger and have higher thyroid hormone levels than solitary nesters. Moreover, we found correlations between corticosterone and glucose, corticosterone and lactate, glucose and lactate, and TG and NEFA levels in arribada but not in solitary nesters. Thus, our combined results suggest that arribada nesters have larger energy reserves and higher metabolic activity than solitary nesters. Our results are consistent with those reported in animals nesting in the Rushikulya Rookery of Orissa, India, where arribada nesters are also larger than solitary nesters [38]. As discussed above, the same individual can nest interchangeably in arribadas or in solitary, but the factors that determine whether an individual joins the arribada or nests in solitary remain unknown [2,4]. Our results suggest that arribadas are energetically more expensive than solitary nesting and that bigger animals with larger energetic reserves join the arribadas. As capital breeders [39–41], sea turtles feed and build their energy reserves prior to migrating, mating and nesting. Hence, if arribada nesting is more energetically costly than solitary nesting, individuals lacking appropriate energy reserves might choose solitary nesting over joining the arribadas despite losing the potential fitness benefits associated with arribada nesting. Solitary nesting requires shorter inter-nesting intervals. Thus, solitary nesting likely results in less time spent away from the feeding grounds [34], potentially allowing solitary nesters to build their energy reserves and join the arribadas later. Therefore, potential differences in resource availability and allocation might ultimately affect whether an individual joins the arribada. According to life history theory, reproduction is a costly life history trait, and our results suggest that nesting in arribada is more energetically expensive than nesting in solitary, though it carries potential fitness benefits such as increased genetic exchange [8].

### Oxidative damage is a potential trade-off of the fitness benefits associated with arribada nesting

The cost of reproduction represents one of the most fundamental life history trade-offs [16], but there is inconclusive evidence about whether oxidative stress is a proximal cost of reproductive investment [14]. Some results support this idea [15–17], while others refute it [18,19]. Increased reproductive effort can promote susceptibility to oxidative stress derived from either exhaustion of antioxidant defenses or dysregulated oxidant generation due to increased metabolic activity, but work with laboratory versus wild animals often yields conflicting results [42]. Moreover, differences in reproductive strategies within wild vertebrates likely result in differential susceptibility to oxidative stress [20–22]. As capital breeders, sea turtles depend on stored energy to fuel reproductive expenditure [43,44]. Whether capital breeders have higher susceptibility to oxidative stress than income breeders remains unknown. In free ranging macaques [23] and North American red squirrels [25] reproduction increases oxidative stress. In contrast, reproduction does not affect lipid peroxidation (measured as thiobarbituric acid reactive substances) in Children’s pythons [45], but in aspic vipers [46] and northern elephant seals, reproduction increases oxidative stress despite concurrent upregulation of antioxidant defenses [24]. Here we found higher lipid peroxidation and protein oxidation levels in arribada compared to solitary nesters. We also found that arribada nesting is potentially more energetically costly than solitary nesting. These results suggest that higher energy expenditure in olive ridley turtles nesting in arribadas is associated with increased oxidative stress. In northern elephant seals, breeding increases lipid peroxidation and oxidative DNA damage in males but not in females [24]. Elephant seals are polygynous, sexually dimorphic capital breeders, with adult males being up to eight times heavier than females [47,48]. Males compete for position in a dominance hierarchy used to control access to females [47–50]. Thus, energy expenditure associated with breeding is higher in male than in female elephant seals [51]. Hence, in capital breeders with unique reproductive behaviours that result in increased energy expenditure, oxidative stress might be a trade-off for potential fitness benefits associated with such behaviours.

In conclusion, we found that arribada nesting in olive ridley sea turtles is associated with increased levels of circulating oxidative damage compared to solitary nesting. Moreover, our results suggest that arribada nesters have higher metabolic activity than solitary nesters. Thus, oxidative stress may be a trade-off for a reproductive mode that carries increased fitness benefits in a capital breeder. Whether these trade-offs are present in other capital breeders remains unknown and undoubtedly warrants further investigation.

## Acknowledgements

We thank Centro Mexicano de la Tortuga and Campamento Tortuguero Palmarito personnel, especially Ernesto Albavera-Padilla for his assistance with sampling logistics and useful discussions, and Allison Raymundo, Carmelo Ambrosio, Alberto Jarquín and Antonio Santiago for their assistance during fieldwork. We thank Cooperativa la Tortuga Feliz, Heradio-Santillán and Sóstenes Rodríguez-Reyes from La Escobilla Campamento Tortuguero, and Dr. Javier Beltran-Robledo for their assistance with fieldwork and sample processing.

## Author’s contributions

Conceptualization: B.G.A and J.P.V.-M. Data curation: B.G.A., D.C.E., and D.E.C. Formal analysis: B.G.A. Investigation: B.G.A., D.C.E., M.H-M., and E.M.L-R. Methodology: B.G.A., D.C.E., H.M-L., E.M.L-R., and D.E.C. Project Administration: J.A.M-V, H.M-L. Resources: M.H-M., E.M.L-R., H.M-L., D.E.C, and J.P.V.M. Supervision: J.P.V.M. Writing – original draft: B.G.A. and J.P.V.M. Writing – review and editing: J.P.V.M. All authors gave final approval for publication and agree to be held accountable for the work performed therein.

